# Identification and characterization of a poly(ε-caprolactone)-degrading enzyme with a unique sequence profile from the marine bacterium *Alloalcanivorax gelatiniphagus*

**DOI:** 10.64898/2026.03.04.709486

**Authors:** Haruno Kusumoto, Shin-ichi Hachisuka, Kyogo Iseki, Hiroshi Kikukawa, Ken’ichiro Matsumoto

**Author notes:** Corresponding author: Shin-ichi Hachisuka.

## Abstract

Poly(ε-caprolactone) (PCL) is a well-known biodegradable polyester and is among the few polyesters susceptible to degradation in marine environments; however, marine-derived PCL-degrading enzymes remain poorly characterized. Here, we searched for PCL-degrading enzymes from the marine bacterium *Alloacanivorax gelatiniphagus* JCM 18425 using a genome-based approach. Five candidate genes were predicted, and one encoded protein, designated Ag0826, was identified as a PCL depolymerase. Recombinant Ag0826 was expressed, purified, and biochemically characterized. The enzyme exhibited optimal activity at 35–40°C and pH 8.0, although it showed limited thermal stability. Substrate specificity was compared with that of leaf-branch compost cutinase (LCC), a well-characterized poly(ethylene terephthalate) (PET) hydrolase, using various polyesters. Both enzymes exhibited largely overlapping substrate ranges with respect to the presence or absence of monomer conversion activity across the tested substrates. Ag0826 slightly degraded PET to terephthalic acid, indicating potential PET-hydrolyzing activity; its conversion rate, however, was substantially lower than that of LCC, suggesting that Ag0826 exhibits a substrate preference differing from LCC. Phylogenetic analysis based on amino acid sequences revealed that Ag0826 formed a separate clade from LCC and *Is*PETase (from *Ideonella sakaiensis*). At a broader level, Ag0826 was positioned near *Halo*PETase1 (from *Halopseudomonas pachastrellae*), which has been proposed as a Type III PET hydrolase; in contrast, residues corresponding to the substrate-binding subsites were similar but not identical between the two enzymes. These results suggest that Ag0826 broadly belongs to the group of known PET hydrolases, yet it exhibits a partially distinct sequence profile even within this enzyme family.

## Introduction

Plastics have become indispensable in modern society due to their low cost, versatility, and durability; consequently, global plastic use has increased to hundreds of millions of tonnes annually [1]. However, the vast majority ultimately ends up as waste in landfills or the natural environment, potentially causing severe ecological impacts and, indirectly, human health problems via the accumulation and ingestion of microplastics [2]. In this context, biodegradable plastics, which can be assimilated by microorganisms, have attracted increasing attention as a promising alternative that may mitigate the environmental burden of plastic pollution [3].

Poly(ε-caprolactone) (PCL), structurally defined as poly(6-hydroxyhexanoic acid) [P(6HHx)], is a highly biodegradable polyester and one of the few plastics susceptible to degradation even in marine environments with low microbial activity [4]. PCL is not a naturally occurring polymer but a chemically synthesized one; therefore, dedicated natural enzymes for its degradation are not expected to exist. Nevertheless, cutinase-like enzymes, probably related to cutinases that hydrolyze the plant polyester cutin, as well as lipase-like enzymes, have been identified as capable of degrading PCL [5], likely owing to the structural similarity between PCL and their natural ester-containing substrates. In addition, leaf-branch compost cutinase (LCC), which is widely recognized as a highly active poly(ethylene terephthalate) (PET) hydrolase, has also been shown to degrade PCL [6]. These findings suggest that the ability of some polyester hydrolases to act on synthetic polymers may stem from their relatively broad substrate specificity.

Numerous bacteria and enzymes capable of degrading PCL have been identified to date [7]. However, reports of PCL-degrading microorganisms and enzymes originating from marine environments remain extremely limited [4]. To the best of our knowledge, only a small number of marine bacterial PCL-degrading enzymes have been characterized so far, including ABO2449 from *Alcanivorax borkumensis* (Q0VLQ1) [8], SM14est from *Streptomyces* sp. SM14 (DAC80635) [9], PET27 from *Aequorivita* sp. CIP111184 (SRX54365) [10], APH*_Hsp_* from *Halopseudomonas* sp. MFKK-1 (WP_228247074) [11], MmCut3 from *Mycobacterium marinum* (ACC39554) [12], and *Halo*PETase1 from *Halopseudomonas pachastrellae* (ONM42822) [13]. Although the number of reported marine-derived PCL-degrading enzymes has increased in recent years, further studies are still required, as elucidation of PCL hydrolysis mechanisms mediated by diverse classes of enzymes may provide valuable insights for the rational design and development of novel biodegradable materials.

In this study, we focused on a marine bacterium, *Alloalcanivorax gelatiniphagus* JCM 18425, with an available genome sequence and demonstrated its ability to degrade PCL. We identified a PCL-degrading enzyme encoded in the genome of the strain and purified the enzyme for biochemical characterization. Sequence analysis indicated that the enzyme shares sequence similarity with PET hydrolases, prompting a comparison with related enzymes to highlight its characteristic motifs. This work provides new insights into the enzymatic mechanisms underlying PCL degradation in marine environments and contributes to a broader understanding of how marine microorganisms participate in the biodegradation of synthetic polyesters.

## Materials and Methods

### Culture media

Unless specifically indicated, reagents were sourced from FUJIFILM Wako Pure Chemical Corporation (Osaka, Japan), Junsei Chemical (Tokyo, Japan), Kanto Chemical Co., Inc. (Tokyo, Japan), Nacalai Tesque (Kyoto, Japan), and Tokyo Chemical Industry (Tokyo, Japan).

Lysogeny broth (LB), Marine Broth 2216 (MB; Becton, Dickinson and Company, Franklin Lakes, NJ, USA), artificial seawater-based (ASWB) [14] media were used in this study. The LB medium contained 10 g/L tryptone, 5 g/L yeast extract, and 10 g/L NaCl. The ASWB medium contained 36 g/L Daigo ASW (Shiotani M.S.; Hyogo, Japan), 1 g/L NH_4_NO_3_, 0.5 g/L yeast extract, 1 mL/L trace metal solution, and 1 mL/L vitamin solution. The components of the trace metal solution and vitamin solution are described in a previous paper [14]. The pH of ASWB was approximately 7.5. Solid plates were prepared by adding agar powder at a final concentration of 1.5% (w/v).

### Bacterial Strains

*A. gelatiniphagus* JCM 18425 [15] was obtained from Japan Collection of Microorganisms (JCM), RIKEN BioResource Research Center (Ibaraki, Japan).

*Escherichia coli* JM109 was used for plasmid cloning, and *E. coli* BL21(DE3) was used for recombinant protein expression.

### Polymer materials

The polymers used in this study are summarized in Table 1. Molecular weight determination by gel permeation chromatography (GPC) was performed as described previously [16].

**Table 1.**
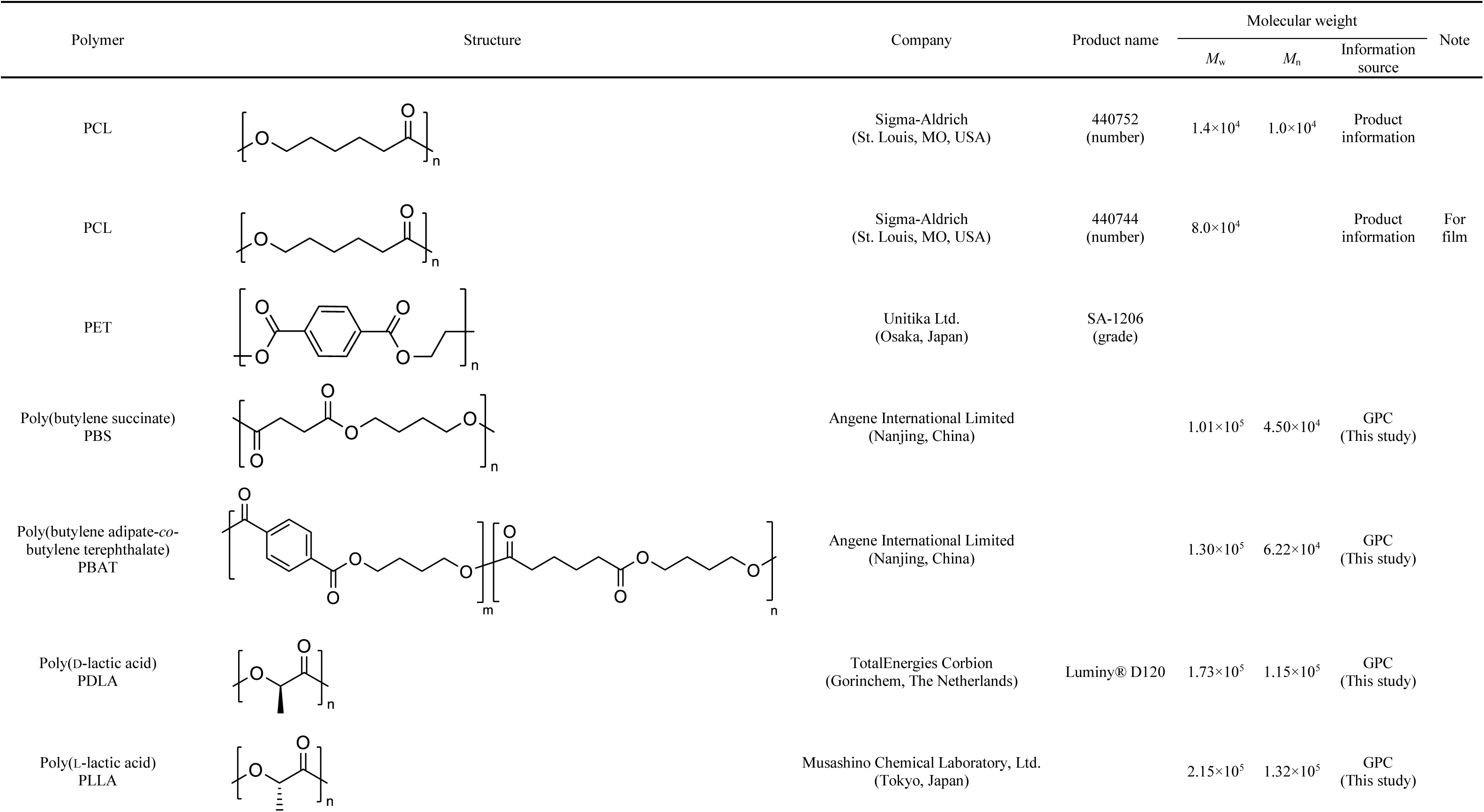
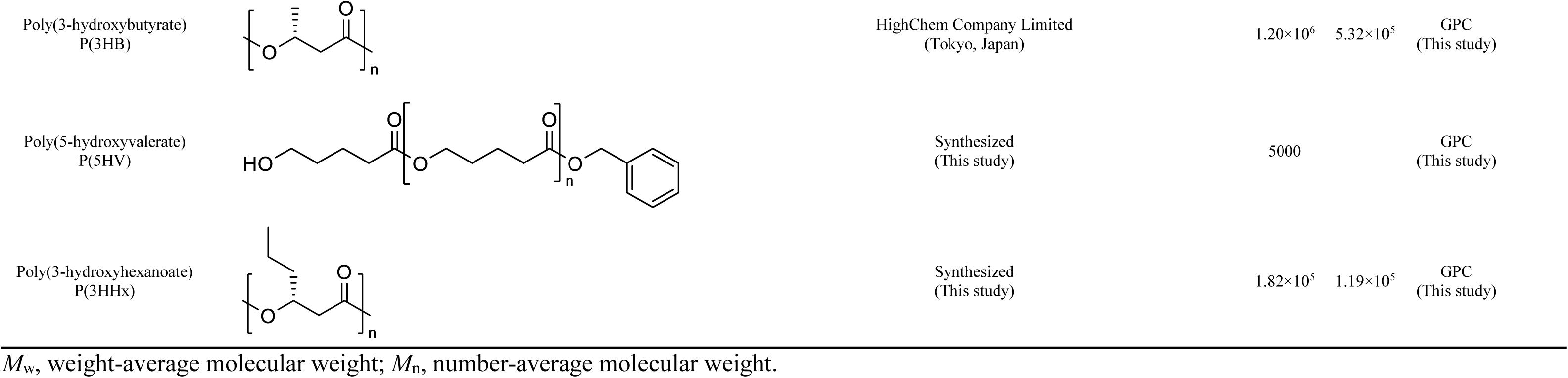
Polymers used in this study.

P(5HV) was synthesized through ring-opening polymerization of δ-valerolactone, as described previously [17]. In brief, δ-valerolactone (5.56 g, 48 mmol, 50 equiv.) and boric acid (0.065 g, 0.960 mmol) were added to a dried 100 mL two-necked round-bottom flask under a nitrogen atmosphere. Benzyl alcohol (0.10 g, 0.960 mmol, 1.0 equiv.) was then added to initiate the polymerization. The reaction was conducted at 130°C for 72 h with vigorous stirring under a nitrogen atmosphere. After completion of the reaction, the products were dissolved in dichloromethane, and a large excess of cold ethanol was added to remove residual monomer and initiator. The purified product was dried under vacuum.

P(3HHx) was biosynthesized using recombinant *E. coli* JM109 expressing the polyhydroxyalkanoate synthase PhaC_AR_ F314H variant, as described previously [18]. The strain was cultivated with 3HHx-Na as the sole monomer precursor. 3HHx-Na was prepared by hydrolysis of ethyl 3-hydroxyhexanoate, as described previously [19].

### Preparation of polymer emulsions

The emulsion used for the medium was prepared as follows. PCL was dissolved in dichloromethane, and ultrapure water was added to achieve a 10 g/L PCL concentration. The mixture was then dispersed using an ultrasonic disperser UD-211 (Tomy Seiko Co., Ltd., Tokyo, Japan) at 20 kHz for 5 min. After allowing the dispersion to stand for approximately 10 min, the supernatant was aseptically added to the medium.

For enzyme assays, PCL emulsion was prepared as follows. PCL was dissolved in dichloromethane, and 0.001% (v/v) Plysurf A210G (DKS Co., Ltd., Kyoto, Japan) and ultrapure water were added to the desired volume. The mixture was then dispersed by sonication at 20 kHz for 5 min using an ultrasonic disruptor UD-211 (Tomy Seiko Co., Ltd.). The resulting dispersion was heated under gentle stirring to completely evaporate the dichloromethane. The resulting emulsion was adjusted to the desired volume with Milli-Q water and used immediately for enzyme assays.

For the substrate specificity analysis, emulsions of PDLA, PLLA, P(3HB), P(3HHx), PBS, PBAT, and P(5HV) were prepared using the same method as above. By contrast, PET emulsion was prepared via solvent displacement, in which 1,1,1,3,3,3-hexafluoro-2-propanol (HFIP) was used as the PET solvent instead of dichloromethane. The method was performed as described previously with minor modifications [20]. Briefly, PET pellets were completely dissolved in HFIP at room temperature to obtain a clear PET/HFIP solution. This solution was then added dropwise into ultrapure water under vigorous stirring to form a milky PET dispersion. After the addition was complete, the mixture was further stirred to promote emulsification. The HFIP was then removed by heating and stirring, and the resulting PET emulsion was adjusted to the desired volume with ultrapure water and used immediately for enzyme assays.

### Degradation assay using medium plates

ASWB containing approximately 2 g/L PCL emulsion was used as the medium for the degradation assay. *A. gelatiniphagus* precultured on an MB containing 1 g/L sodium acetate plate was inoculated into the ASWB-PCL using a sterile toothpick. The plate was incubated at 30°C for a week.

### Protein nomenclature

Protein identifiers used in this study were defined based on the submitted GenBank assembly of *A. gelatiniphagus* JCM 18425 (accession GCA_005938655.1). In this assembly, coding sequences (CDSs) were annotated with locus tags ranging from GCA_005938655.1_00001 to GCA_005938655.1_03817. For convenience, each protein was assigned a simplified identifier by combining the initials of the species name (*A. gelatiniphagus*, “Ag”) with the last four digits of the corresponding locus tag (i.e., GCA_005938655.1_0XXXX). Thus, proteins were referred to in this study as AgXXXX.

### Plasmids

The genomic DNA was extracted using the Wizard Genomic DNA Purification Kit (Promega; Madison, WI, USA). The target genes, excluding the signal sequences, were amplified by PCR from the extracted genomic DNA using primer sets (Table S1) and inserted into pET-21a(+) at the NdeI–XhoI sites. These plasmids were designated as pET-21a-AgXXXX.

For the analysis of the purified enzyme, Ag0826 gene excluding the signal sequence was codon-optimized for *E. coli* to enhance its expression level. The reconstituted gene sequence was synthesized by Twist Bioscience (South San Francisco, CA, USA) and the sequence are shown in Fig. S1. Ag0826 codon-optimized gene [Ag0826(opt)], was amplified by PCR using the primer set (Table S1) and inserted into the NdeI and XhoI sites of the plasmid pET-21a(+).

The LCC expression plasmid was constructed by reverting the sequence of a previously reported ICCG mutant LCC expression plasmid [16] to the original LCC sequence, with codon selection guided by the previous study [21]. Although pET-21b(+) was used as the backbone, the LCC gene was inserted between the NdeI and XhoI sites; therefore, the resulting plasmid sequence was identical to that obtained using pET-21a(+).

### Enzyme preparation

The *E. coli* BL21(DE3) cells were transformed with each of the pET-21a-AgXXXX plasmids. The transformant was cultured in LB supplemented with 100 µg/L ampicillin sodium (Amp-Na) at 37°C, and isopropyl β-D-1-thiogalactopyranoside (IPTG; final concentration, 0.1 mM) was added to the medium when the OD_600_ reached 0.6-0.7. Then, the cells were further cultivation at 16°C for 24 h. Cells ware harvested by centrifugation (5,000 × g, 10 min, 4°C), resuspended in 50 mM Tris-HCl buffer (pH8.0) with 0.15 M NaCl, and disrupted by sonication (20 kHz) using an ultrasonic disruptor UD-211 while being chilled in ice water. The resulting cell lysate was centrifuged, and the supernatant was used as the cell-free extract for the first screening. The total protein concentration was determined using the Bradford method.

For detailed analysis of the Ag0826 protein, *E. coli* BL21(DE3) cells were transformed with plasmid pET-21a-Ag0826(opt). The transformant was cultured in the same manner as described above. Cells were harvested by centrifugation (5,000 × g, 10 min, 4°C), resuspended in 50 mM Tris-HCl buffer (pH8.0) with 0.3 M NaCl and 5 mM imidazole, and disrupted by sonication (20 kHz) while being chilled in ice water. Subsequently, the cell-free extract was obtained from the 3L culture was subjected to 0.5 mL of TALON affinity resin (Takara Bio., Shiga, Japan). Proteins mainly containing the His_6_-tagged Ag0826 were eluted with 50 mM Tris-HCl buffer (pH8.0) containing 0.3 M NaCl and 200 mM imidazole. The eluted fraction was subjected to a PD-10 desalting column (Cytiva, Marlborough, MA, USA). The concentration of the purified protein was determined by the absorption value at 280 nm. The molar extinction coefficient of the protein was calculated using the ExPASy ProtParam tool (https://web.expasy.org/protparam/).

The LCC was prepared using Terrific Broth (Becton, Dickinson and Company) with a reduced culture volume, as high expression levels were obtained, and was otherwise prepared in the same manner as Ag0826.

### First screening of PCL-degrading activity

The reaction mixture contained 100 mM potassium phosphate buffer (KPB, pH8.0), 2.6 g/L PCL emulsion, 1.0 mg/mL AgXXXX-expressing cell-free extract. The mixture was incubated at 30°C for 2 h, after which the supernatant was collected by centrifugation. The degradation of PCL was evaluated by quantifying 6HHx, the monomer unit of PCL, via high-performance liquid chromatography (HPLC).

### HPLC analysis

The released degradation products were analyzed by HPLC using an LC-4000 series system (JASCO Corporation, Tokyo, Japan) under the following conditions. For sample preparation, 50 μL of 1 M H_2_SO_4_ was added to 1 mL of the reaction mixture. After vortexing, the mixture was incubated on ice for 10 min and centrifuged. The supernatant was filtered through a 0.22 μm membrane filter and used for HPLC analysis.

An Aminex HPX-87H column (300 × 7.8 mm I.D.; Bio-Rad Laboratories, Hercules, CA, USA) was used for the detection of 6HHx, 5HV, LA, 3HB, 3HHx, succinic acid, 1,4-butanediol, adipic acid, ethylene glycol, terephthalic acid (TPA), mono(2-hydroxyethyl) terephthalate (MHET), and bis(2-hydroxyethyl) terephthalate (BHET). MHET and BHET were detected but not quantified. The column temperature was maintained at 60°C. The mobile phase consisted of 7 mM H_2_SO_4_ (isocratic elution) at a flow rate of 0.6 mL/min. For PET and PBAT degradation products, including TPA, the flow rate was adjusted to 0.8 mL/min. The eluted compounds were detected by UV absorption at 210 or 240 nm; 1,4-butanediol and ethylene glycol were detected using a refractive index detector. Peaks were identified and quantified using external calibration curves prepared from authentic standards.

6HHx-Na and 5HV-Na, used as standards, were synthesized through ring-opening ε-caprolactone or δ-valerolactone with NaOH as described previously [22]. The identity and purity of the products were confirmed by ^1^H NMR.

### Effects of temperature, pH, metal ions, and NaCl on the PCL-degrading activity

The reaction mixture consisted of 100 mM KPB (pH 8.0) and 0.5 g/L PCL emulsion. After pre-incubation at 35°C for approximately 5 min, the reactions were initiated by adding purified enzyme (final concentration, 2.5 µg/mL). Enzyme activity was measured by monitoring the decrease in OD_650_ at 35°C for 5 min using 1-cm light-path cuvettes. Values immediately after the start of the measurement were not used for the calculation due to instability. Unless otherwise stated, assays were performed under the above conditions, with individual parameters varied as described below. Enzyme activity was determined from the rate of decrease in OD_650_, and relative activities were calculated by normalizing the observed rates to a reference value defined for each experiment.

To investigate the effect of temperature on enzyme activity, assays were performed over a temperature range of 10–60°C. To examine the residual activity after thermal treatment, the enzyme was incubated at various temperatures (10–40°C) for 30 min, followed by activity measurement at 35°C for 5 min. The optimal pH was determined by assaying the enzyme in 100 mM buffers of various pH values: sodium citrate buffer (pH 4–6), KPB (pH 6–8), and Tris-HCl buffer (pH 8–9). To examine the effects of metal ions, various metal salts (K^+^, Ca^2+^, Mg^2+^, Mn^2+^, and Na^+^) or ethylenediaminetetraacetic acid (EDTA) were added to the reaction mixture containing the enzyme in the absence of substrate at final concentrations of 1 or 10 mM. The mixtures were incubated on ice for 30 min, and the reactions were then initiated by the addition of substrate under the standard assay conditions (35°C). To investigate the effect of NaCl concentration on enzyme activity, NaCl was added to the reaction mixture at final concentrations ranging from 0 to 1.0 M.

### PCL film degradation assay

PCL films were prepared using the solvent casting method. PCL (*M*_n_ = 80,000) was dissolved in CHCl_3_, then cast onto a dish while being filtered through a 0.22 μm PTFE filter and dried at room temperature for 3 days. Individual PCL films (1.0 × 0.5 cm) were incubated with purified Ag0826 (final concentration, 10 or 30 µg/mL) in 100 mM KPB (pH 8.0) at 35°C for 24 h. The surface morphology of PCL films before and after enzymatic hydrolysis was examined using a JEOL JSM-6510LA SEM-EDS (JEOL Ltd., Tokyo, Japan) operated at an accelerating voltage of 10 kV. Prior to analysis, the film surfaces were coated with platinum by sputtering.

### Examination of substrate specificity

To determine the substrate specificity of Ag0826, PDLA, PLLA, P(3HB), P(3HHx), PBS, PBAT, P(5HV), and PET were used as substrates in addition to PCL. LCC was used as a reference enzyme for comparison of polyester-degrading activities. Polymer degradation was evaluated based on the amount of monomers produced, as determined by HPLC. The reaction mixture consisted of 100 mM KPB (pH 8.0), 2 g/L polymer emulsion, and 30 µg/mL purified enzyme. Reactions were incubated at 25°C for 24 h and quenched by the addition of H_2_SO_4_. After centrifugation, the resulting supernatants were analyzed by HPLC.

### Phylogenetic analysis

A phylogenetic tree was constructed using amino acid sequences of previously reported depolymerases that degrade polyesters such as PCL and PET, together with homologous sequences of the target proteins. To avoid over-representation of closely related proteins, homologous sequences from the same genus as the target proteins were excluded. Phylogenetic trees were inferred by the maximum likelihood method implemented in MEGA12 [23], and branch support was evaluated by bootstrap analysis with 100 replicates.

### Multiple sequence alignment analysis

Multiple sequence alignment was generated using CLUSTAL W [24], and the alignment was visualized and annotated with ESPript version 3.2 [25].

## Results

### Characterization of PCL-degrading activity in *A. gelatiniphagus* JCM 18425

*A. gelatiniphagus* JCM 18425 was cultured on a PCL dispersion plate. A clear zone was observed around the bacterial growth after 7 days of incubation at 30°C (Fig. 1). The formation of this clear zone indicates degradation of PCL in the medium surrounding the growth area, suggesting that the strain produces extracellular PCL-degrading enzyme(s).

**Fig. 1.**
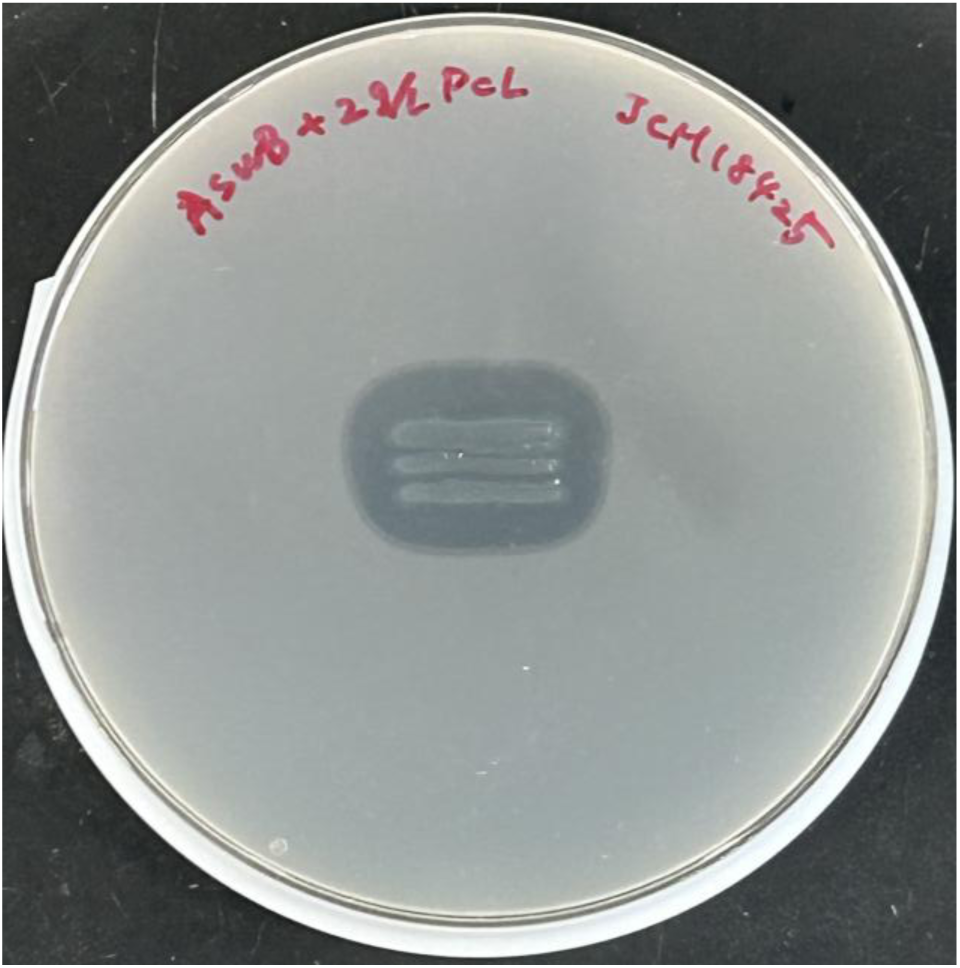
PCL degradation assay of *A. gelatiniphagus* JCM 18425 using medium plate. The strain was cultured on an ASWB-PCL (2 g/L) medium plate at 30°C for 7 days.

### Screening for PCL-degrading enzyme(s)

To predict PCL-degrading enzymes, the genome sequence of *A. gelatiniphagus* JCM 18425 was examined using the EzBioCloud platform [26]. Proteins annotated as lipases or cutinases, as well as those showing sequence similarity to previously reported microbial PCL-degrading enzymes, including CAA83122 [27, 28], Q99174 [29], AEV21261 (LCC; [6]), and LC189557 [30], were selected. Among these candidates, proteins predicted to possess a secretion signal peptide (Sec/SPI) were further screened using SignalP 6.0 [31]. As a result, proteins Ag1351, Ag1433, Ag1855, Ag2147, and Ag0826 were selected (Table 2).

**Table 2.**
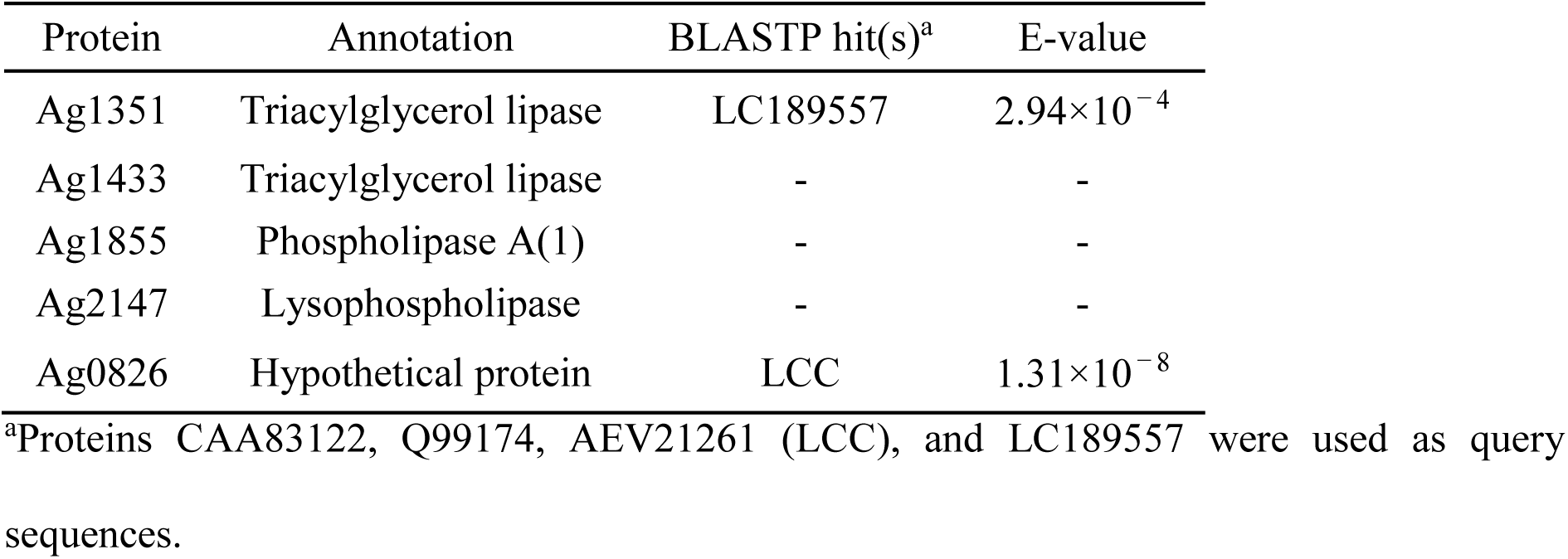
Selected PCL-degrading enzyme candidates.

| Protein | Annotation | BLASTP hit(s)<sup>a</sup> | E-value |
| --- | --- | --- | --- |
| Ag1351 | Triacylglycerol lipase | LC189557 | $2.94 \times 10^{-4}$ |
| Ag1433 | Triacylglycerol lipase | - | - |
| Ag1855 | Phospholipase A(1) | - | - |
| Ag2147 | Lysophospholipase | - | - |
| Ag0826 | Hypothetical protein | LCC | $1.31 \times 10^{-8}$ |

<sup>a</sup>Proteins CAA83122, Q99174, AEV21261 (LCC), and LC189557 were used as query sequences.

To examine the PCL-degrading activity of the five selected candidates, the regions lacking the signal sequences of each protein were expressed in *E. coli*. Protein expression was analyzed by SDS-PAGE using cell-free extracts (Fig. S2). Strong expression at the expected molecular weight was observed for Ag2147 and Ag1855, whereas clear expression was not detected for the other candidates.

Screening for PCL-degrading activity was performed using the five candidate proteins. Reaction mixtures containing PCL emulsion and cell-free extracts expressing each candidate were incubated, and the formation of the monomer 6HHx was analyzed by HPLC. Among the tested candidates, a distinct product peak at a retention time corresponding to 6HHx was detected in the reaction mixture containing the Ag0826-expressing cell-free extract (Fig. 2), whereas no such peak was detected in the negative controls (empty vector control or no cell-free extract). In contrast, reaction mixtures containing cell-free extracts expressing the other candidates exhibited only baseline signals comparable to those of the negative controls. These results indicate that Ag0826 possesses PCL-degrading activity.

**Fig. 2.**
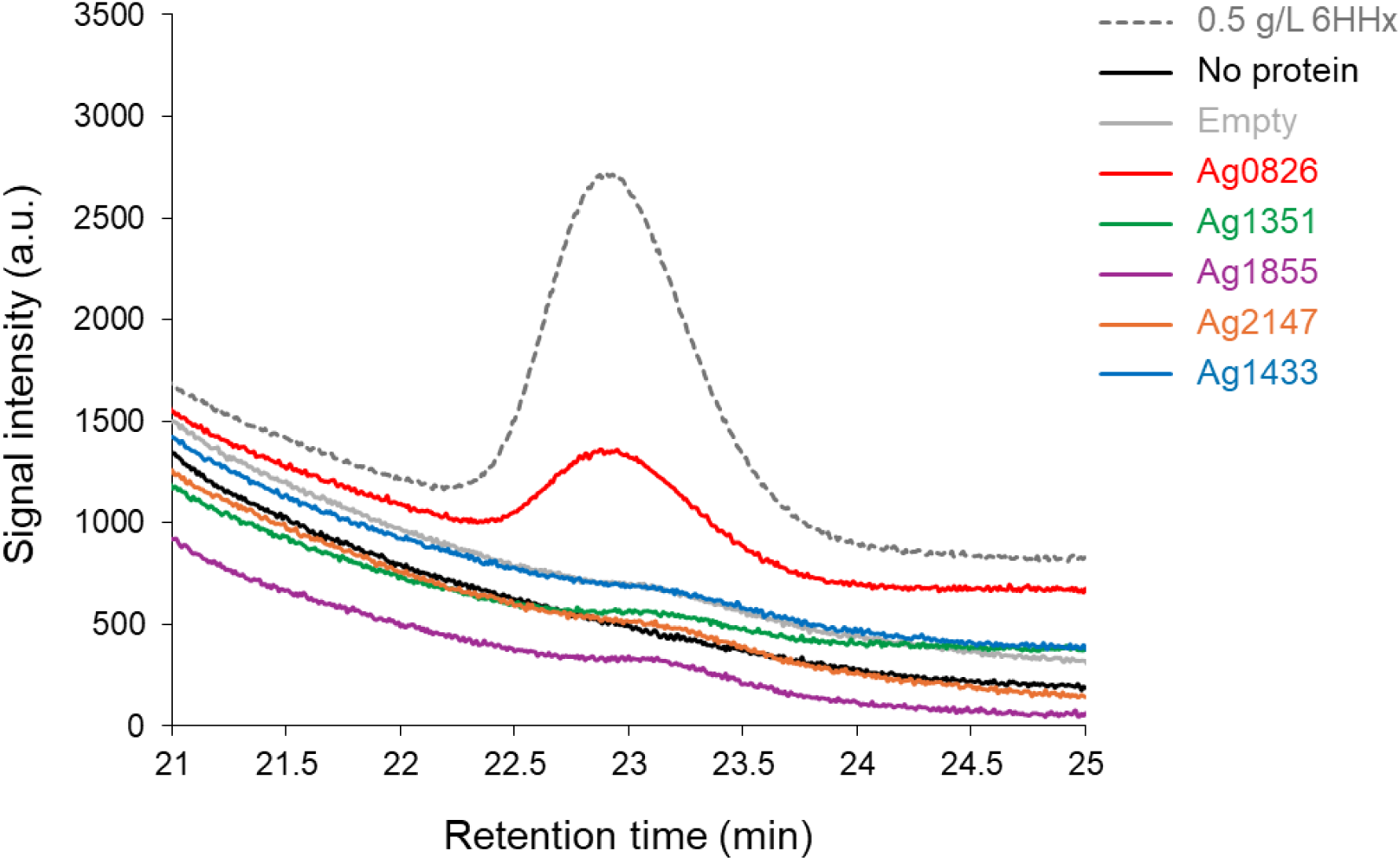
HPLC chromatograms of soluble fractions after reaction of PCL emulsion with cell-free extracts expressing each candidate protein. The retention time of 6HHx was approximately 22.9 min. “No protein” and “Empty” indicate reaction mixtures without cell-free extract and with cell-free extract from cells harboring the pET-21a(+) empty vector, respectively.

### Biochemical properties of purified Ag0826

Based on the observed PCL-degrading activity of Ag0826, the protein was purified by Co^2+^ affinity chromatography via a C-terminal His_6_-tag for further characterization. SDS-PAGE analysis of the purified fraction showed a major band at approximately 27 kDa, indicating high purity (Fig. S3).

Using purified Ag0826, the effects of temperature, pH, metal ions, and NaCl concentration on PCL hydrolysis were investigated. Enzyme activity was determined by measuring the decrease in turbidity of a PCL emulsion over 5 min, and specific activity was calculated. For comparison among conditions, activities are presented as relative values. Ag0826 exhibited maximal activity at 35–40°C (Fig. 3A). However, residual activity after 30 min of heat treatment began to decrease at temperatures above 20°C and dropped to nearly 0% following treatment at 35–40°C (Fig. 3B). The optimal pH for activity was pH 8.0 (Fig. 3C). Regarding the effects of metal ions (Table 3), the addition of 10 mM Mn^2+^ significantly reduced enzyme activity, although the reason for this inhibition is unclear. In contrast, the addition of other metal ions at concentrations up to 10 mM did not cause marked changes in activity. Furthermore, the activity was not substantially decreased by the addition of EDTA. Taken together, these results suggest that Ag0826 does not require a metal ion as an essential cofactor. Since this enzyme originates from a marine microorganism, the effect of higher NaCl concentrations on activity was also examined (Fig. 3D). Maximum activity was observed at 0.2 M NaCl, and the activity remained higher than that in the absence of NaCl at concentrations up to 0.8 M.

**Fig. 3.**
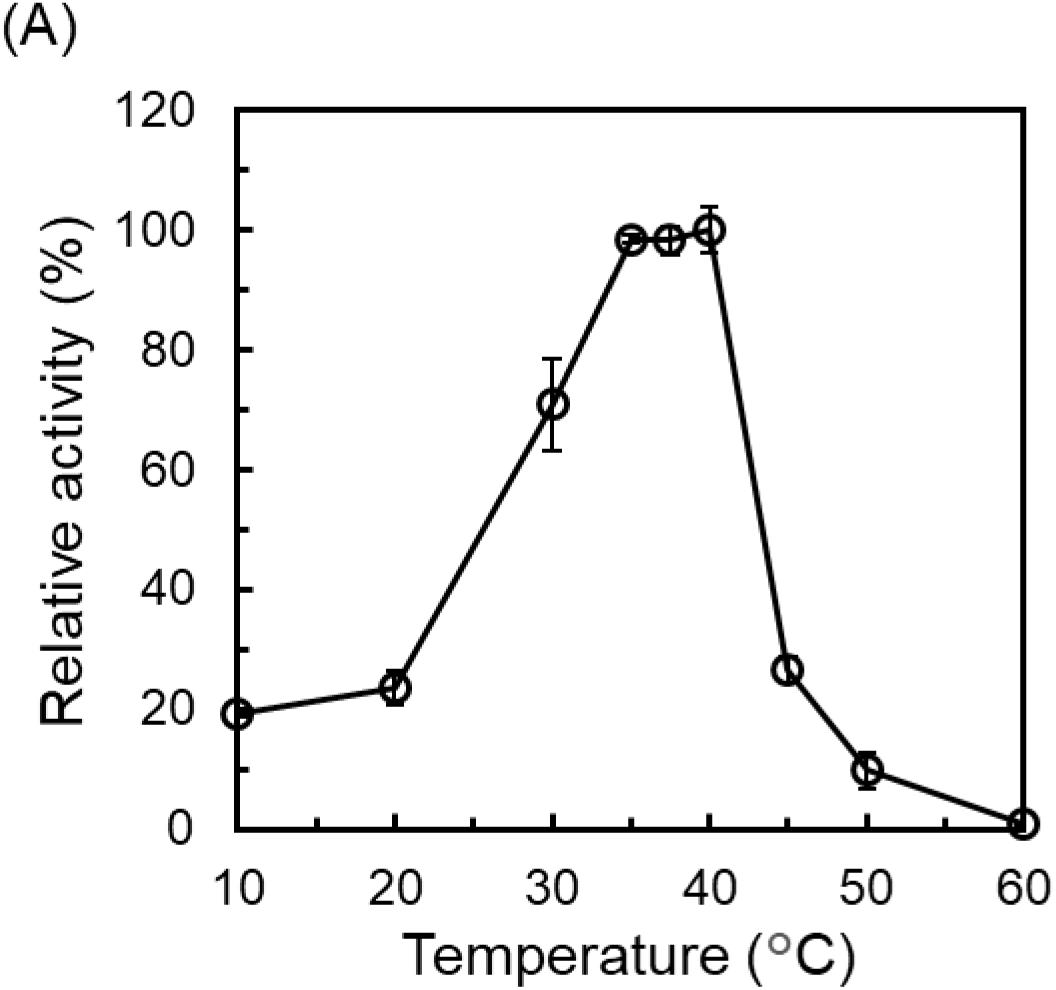
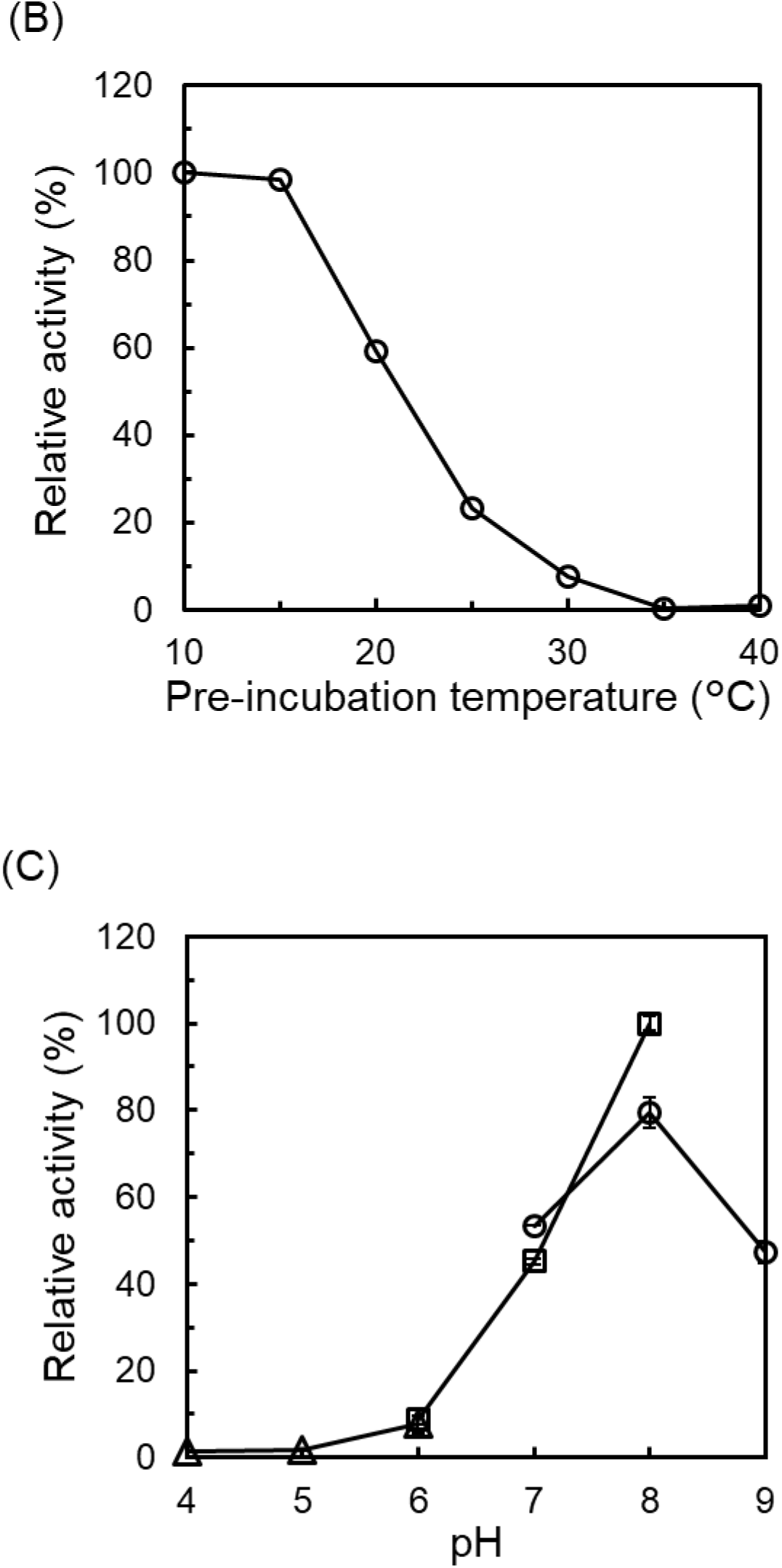
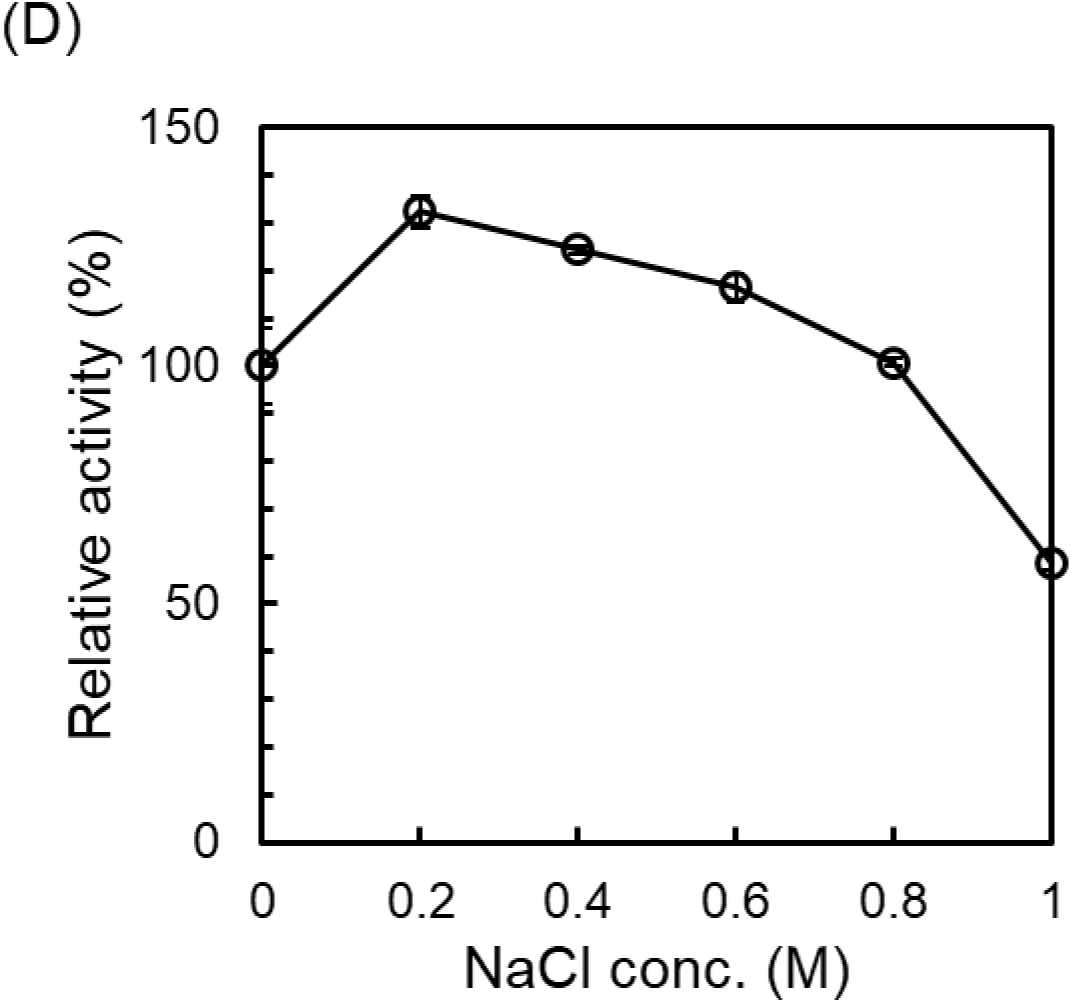
Effects of temperature, pH, and NaCl concentration on the PCL-hydrolyzing activity of purified Ag0826. (A) Temperature dependence of enzyme activity. Reactions were carried out in KPB (pH 8.0) for 5 min. Activity at 40°C was defined as 100%. (B) Thermal stability of Ag0826. Residual activity was measured after pre-incubation at the indicated temperatures for 30 min. The enzymatic reaction was subsequently performed at 35°C for 5 min in KPB (pH 8.0). Activity after pre-incubation at 10°C was defined as 100%. (C) pH dependence of enzyme activity. Reactions were performed at 35°C for 5 min using the following buffers: sodium citrate buffer (triangles), KPB (squares), and Tris-HCl buffer (circles). Activity in KPB (pH 8.0) was defined as 100%. (D) Effect of NaCl concentration on enzyme activity. Reactions were conducted in KPB (pH 8.0) at 35°C for 5 min. Activity in the absence of NaCl (0 M) was defined as 100%. Data in panel (B) were obtained from a single experiment (n = 1), whereas all other data represent the mean of three independent experiments (n = 3). Error bars indicate the standard error of the mean (SEM).

**Table 3.** Effects of metal ions on the PCL-hydrolyzing activity of purified Ag0826.

| Metal ions | Relative activity (%) |  |
| --- | --- | --- |
|  | 1 mM | 10 mM |
| KCl | 89.7 ± 2.2 | 97.0 ± 3.2 |
| CaCl<sub>2</sub> | 100.4 ± 4.6 | 100.6 ± 9.4 |
| MgCl<sub>2</sub> | 103.0 ± 8.5 | 110.6 ± 3.2 |
| MnCl<sub>2</sub> | 84.6 ± 2.4 | 12.4 ± 4.1 |
| NaCl | 93.6 ± 3.0 | 91.6 ± 1.7 |
| EDTA | 83.2 ± 4.9 | 84.3 ± 2.2 |

### PCL film degradation using Ag0826

To examine whether Ag0826 can degrade PCL in film form, PCL films were incubated with purified Ag0826 in KPB (pH 8.0) at 35°C for 24 h, and the film surfaces were subsequently observed by SEM. In the absence of the enzyme, no noticeable surface irregularities were observed (Fig. 4A). In contrast, enzyme-treated films exhibited clear surface roughness, which was attributed to enzymatic degradation (Figs. 4B and 4C). Moreover, the extent of surface erosion appeared to increase with increasing amounts of enzyme added (Figs. 4B and 4C). These results demonstrate that Ag0826 is capable of degrading PCL in film form and can also degrade high-molecular-weight PCL (*M*_w_ = 80,000), whereas the PCL used in the emulsion assays had a lower molecular weight (*M*_w_ = 14,000).

**Fig. 4.**
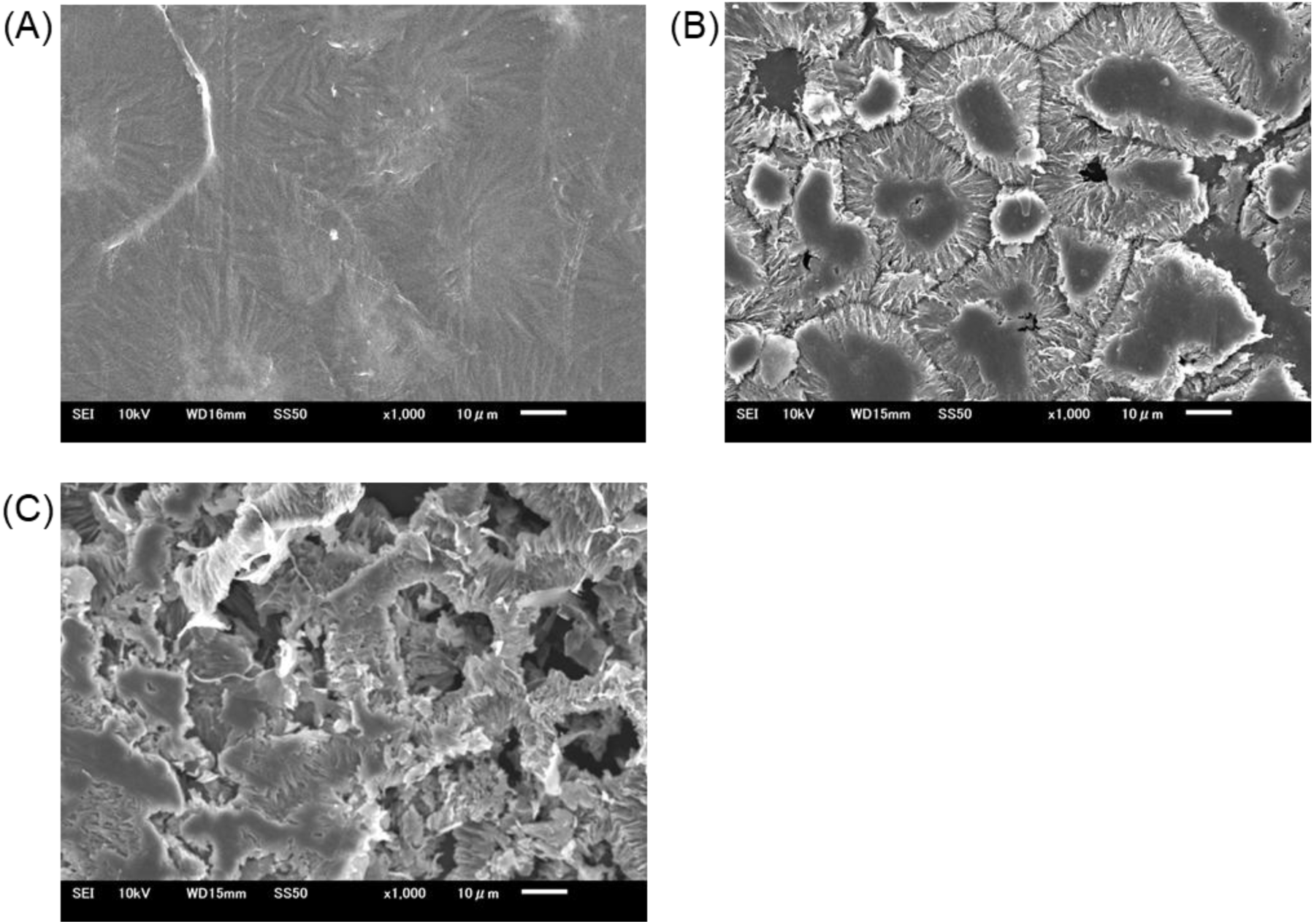
SEM images of PCL films degraded by Ag0826. The magnification was set to 1,000×. The white bar indicates a length of 10 µm. (A) Control condition without enzyme. (B) Condition with 10 µg of enzyme added. (C) Condition with 30 µg of enzyme added.

### Substrate specificity of Ag0826

To investigate hydrolytic activity toward substrates other than PCL, enzymatic assays were performed using emulsions of various polymers. Because of concerns regarding the thermal stability of the enzyme, reactions with PCL were conducted at different temperatures for 24 h (Fig. S4). Monomer production was maximal at 25°C; therefore, this temperature was used for the following substrate specificity assays. LCC was used as a reference enzyme for comparison. After incubation for 24 h, reaction supernatants were analyzed by HPLC to quantify released monomers and to calculate monomer conversion rates (Fig. 5). Among the substrates tested, Ag0826 exhibited clear activity not only toward PCL but also toward P(5HV), PDLA, PLLA, P(3HHx), PBS, and PBAT. Ag0826 produced slightly higher amounts of monomers from PDLA and P(3HHx), whereas LCC showed somewhat higher activity toward P(5HV), PLLA, PBS, and PBAT. Both enzymes exhibited largely overlapping substrate ranges, indicating similarities in their substrate specificities. In contrast, no 3HB generation was detected from P(3HB), whereas LCC generated detectable amounts of 3HB from this substrate. With respect to PET degradation, TPA was detected at very low levels, with a monomer conversion rate below 0.1%. However, the intermediate hydrolysis products MHET and BHET were also detected (Fig. S5), indicating that PET hydrolysis had occurred. By contrast, LCC efficiently converted PET to TPA. Ethylene glycol was also clearly detected in the LCC reaction and was included in the calculation of the monomer conversion rate, whereas it was not reliably detected in the Ag0826 reaction, likely due to the limited sensitivity of the RI detector.

**Fig. 5.**
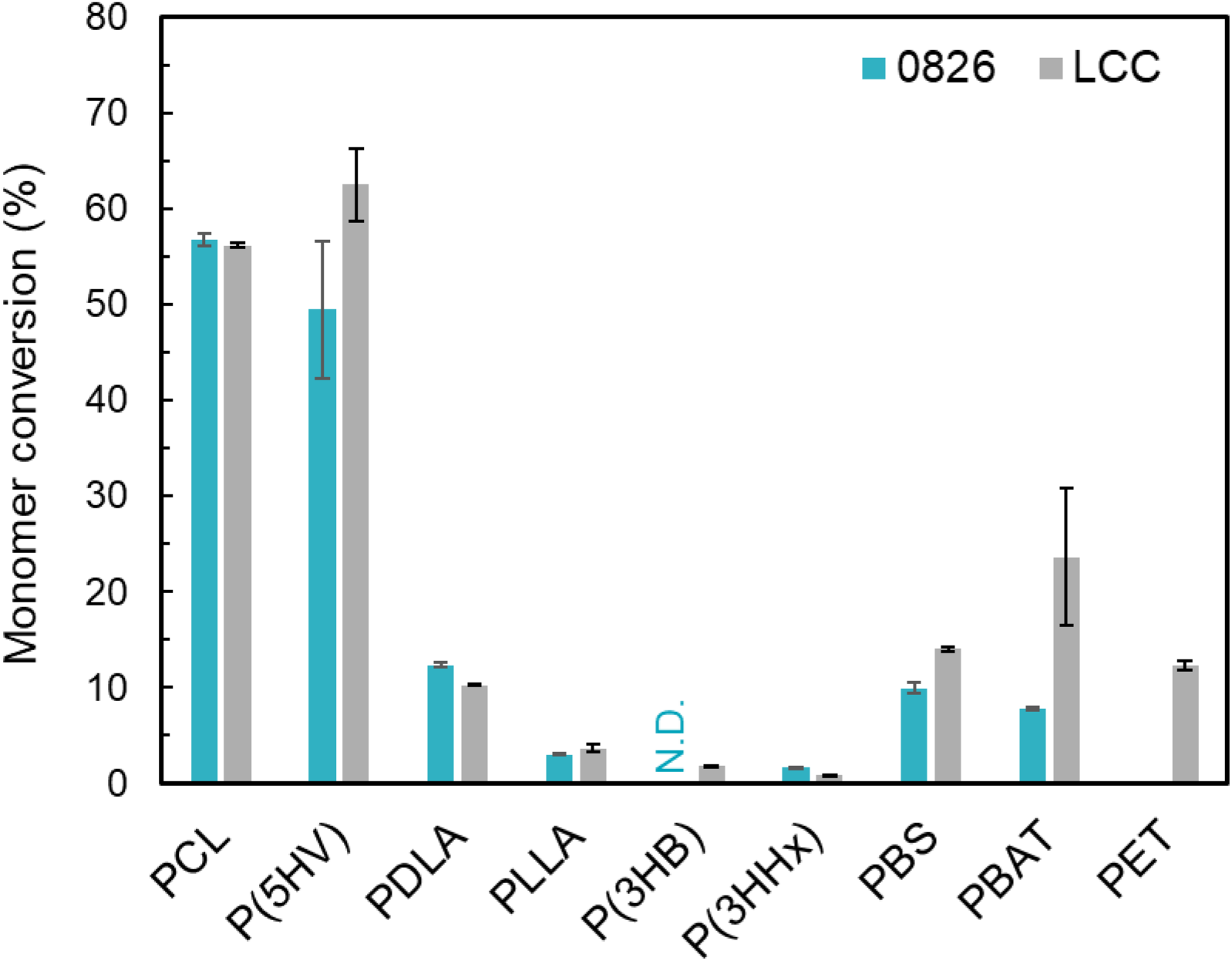
Substrate specificity of Ag0826 and LCC. Monomer conversion (%) after incubation at 25°C and pH 8.0 (100 mM KPB) for 24 h is shown. Monomer conversion (%) was defined as the ratio of the amount of monomer released to the theoretical amount obtainable upon complete hydrolysis of the added polymer emulsion. Monomers released from each polymer were quantified as follows: 6HHx from PCL; 5HV from P(5HV); LA from PDLA and PLLA; 3HB from P(3HB); 3HHx from P(3HHx); succinic acid and 1,4-butanediol from PBS; TPA, 1,4-butanediol, and adipic acid from PBAT; and TPA and ethylene glycol from PET. “N.D.” indicates that the monomer was not detected. Data represent the mean ± SEM of three independent experiments (n = 3).

### Phylogenetic analysis of Ag0826 and related PCL- and PET-degrading enzymes

To clarify the phylogenetic position of Ag0826 among known polyester-degrading enzymes, a phylogenetic tree was constructed using previously reported amino acid sequences of PCL- and PET-degrading enzymes together with sequences homologous to Ag0826. The resulting phylogenetic tree (Fig. 6) was divided into several major clades. The clade containing LCC and the clade containing *Is*PETase corresponded to the previously reported PET hydrolase type I and type II groups, respectively [32, 33]. These types are distinguished based on structural and sequence features, including the presence and position of disulfide bonds, the amino acid composition of the active-site subsites (subsites I and II), and the presence of an elongation loop adjacent to the active site. More recently, in addition to these groups, Turak et al. proposed a type III classification for *Halo*PETase1 and its close relative *Pm*C [13]. Ag0826 was positioned in proximity to these enzymes in the phylogenetic tree, suggesting that it may broadly fall within the type III group. However, Ag0826 shares 51% and 47% sequence identity with *Halo*PETase1 and *Pm*C, respectively, which are slightly lower than the sequence identity between *Halo*PETase1 and *Pm*C (59%). Therefore, further careful analysis is required to definitively assign Ag0826 to the type III classification.

**Fig. 6.**
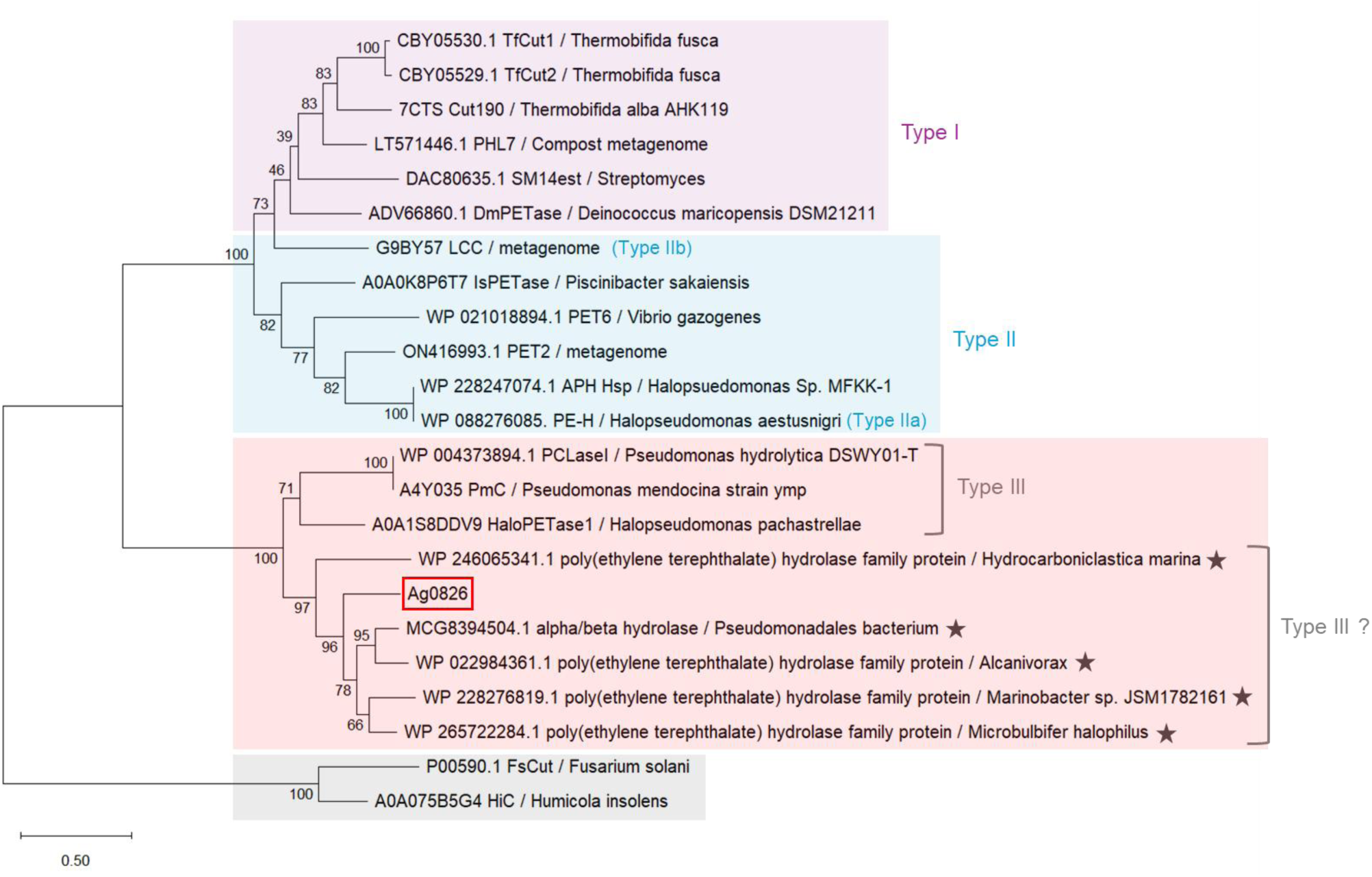
Maximum-likelihood phylogenetic tree of Ag0826 and related enzymes, including known PCL- and PET-hydrolases. Bootstrap analysis was performed with 100 iterations. For PET hydrolase classification, type I enzymes are shown in purple, type II in light blue, and enzymes tentatively assigned to type III in red. The gray clade represents fungal enzymes. Black stars indicate sequences homologous to Ag0826 that were selected for this phylogenetic analysis, whereas the remaining sequences correspond to previously reported PCL- and PET-hydrolases.

## Discussion

In this study, we identified a PCL-degrading enzyme, Ag0826, from the marine bacterium *A. gelatiniphagus* JCM 18425 and characterized its fundamental biochemical properties using the purified recombinant protein. Comparison of its substrate specificity with that of LCC, a well-known PET hydrolase, revealed clear differences in activity toward PET and P(3HB). In contrast, for other tested polyesters, the differences in activity were more moderate, suggesting that while Ag0826 shares some substrate preferences with LCC, it exhibits a distinct catalytic profile. Phylogenetic analysis based on amino acid sequences showed that Ag0826 forms a clade distinct from canonical PET hydrolases such as LCC and *Is*PETase. At a broader evolutionary scale, however, Ag0826 was positioned relatively close to *Halo*PETase1, which has been proposed as a Type III PET hydrolase. Although Ag0826 clustered near *Halo*PETase1, it appeared not to belong to the same subgroup but instead to occupy a slightly different phylogenetic position.

From genome mining, five candidate genes were selected as putative PCL-degrading enzymes, of which only one, Ag0826, exhibited detectable PCL-degrading activity under the conditions tested. However, this result does not allow us to conclude that the remaining candidates lack PCL-degrading activity. Ag2147 and Ag1855 were expressed in soluble forms (Fig. S2), yet no activity was detected in these recombinant proteins, indicating that they were inactive at least under the present expression and assay conditions. Notably, previous studies have reported that changing affinity tags alone can alter the apparent activity of enzymes [13], suggesting that these proteins may not have been obtained in an active form. In contrast, Ag1433 and Ag1351 were not detected as soluble proteins by SDS-PAGE, making it unclear whether they were properly expressed. To determine whether additional PCL-degrading enzymes exist in *A. gelatiniphagus*, future work will require the development of a genetic disruption system for this organism, including targeted deletion of the Ag0826 gene to evaluate its contribution to PCL degradation in vivo.

Ag0826 exhibited optimal activity at 35–40°C and pH 8.0, whereas its residual activity decreased after heat treatment at temperatures above 20°C (Fig. 3), indicating relatively limited thermal stability. These optima are consistent with the reported growth conditions of *A. gelatiniphagus*, which grows best at 37–40°C and pH 7.0–8.0 [15], suggesting that the catalytic properties of the enzyme are well adapted to the physiological environment of the host. The apparent thermal lability of the enzyme, however, may seem counterintuitive for an enzyme with an optimum near 40°C. Nevertheless, because *A. gelatiniphagus* is a marine bacterium, it is likely to function at temperatures lower than its optimal growth temperature, and thus reduced thermal stability may not be a major disadvantage in its natural habitat. Why this enzyme has evolved such a balance between catalytic efficiency and stability remains unclear and warrants further investigation.

Based on the phylogenetic tree, Ag0826 was broadly suggested to be classified as a Type III PETase (Fig. 6). To further examine this classification at the sequence level, we performed a detailed sequence analysis. Multiple sequence alignment was conducted, and amino acid residues involved in disulfide bond formation were inspected (Fig. 7). It is known that Type I PETases possess one disulfide bond at a conserved position, Type II PETases possess two at conserved positions, and Type III PETases possess two disulfide bonds at positions distinct from those of Type I and II enzymes [13, 32, 34]. In Ag0826, two disulfide bonds were observed at positions identical to those in *Halo*PETase1. This structural feature supports the classification of Ag0826 as a Type III PETase.

**Fig. 7.**
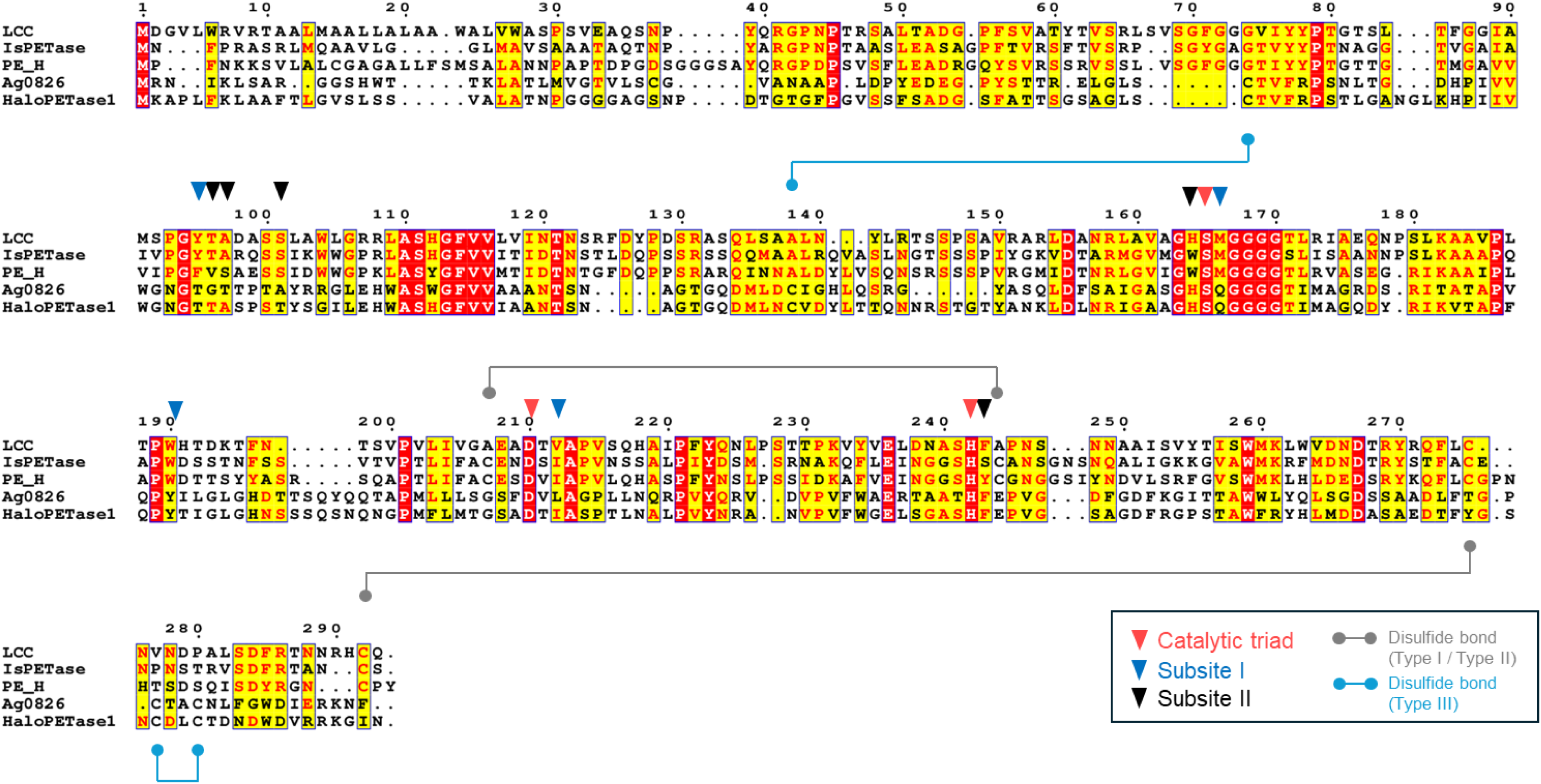
Multiple sequence alignment of Ag0826 with representative PET hydrolases showing catalytic, subsite, and disulfide-bond residues.

Based on sequence alignment, we compared amino acid residues involved in substrate-binding subsites (Table 4). In PETase-family enzymes, subsite I, located adjacent to the catalytic Ser-His-Asp triad, forms a π-stacking clamp with the TPA ring and is generally conserved, whereas subsite II is more variable and influences substrate chain accommodation [32, 34]. In subsite I, Ag0826 differed markedly from typical Type I and Type II PETases; for example, Y95, M166, and W190 of LCC corresponded to T77, Q140, and Y163 in Ag0826. However, this region showed the highest similarity to *Halo*PETase1, suggesting related substrate-binding modes. In subsite II, Ag0826 also differed from previously characterized PETases, although partial similarity to *Halo*PETase1 was retained. These findings imply that even within Type III PETases, further subdivision based on subsite architecture may be possible. Notably, despite substantial sequence differences between Ag0826 and LCC, including in subsite regions, both enzymes exhibited broad substrate specificity and similar patterns regarding the presence or absence of monomer conversion activity for most tested polyesters other than P(3HB) (Fig. 5). This suggests that substrate specificity is not governed solely by subsite residues but also by broader structural factors, such as binding pocket geometry and surface hydrophobicity. Further investigation of these features may provide new targets for engineering enzyme activity and substrate specificity. In addition, the unique sequence–function relationship observed in Ag0826 may offer important insights into the evolutionary trajectory by which PET hydrolases have diversified their substrate recognition and catalytic properties.

**Table 4.** Comparison of subsite residues between Ag0826 and representative PET hydrolases.

| Type | Enzyme | Subsite I |  |  |  | Subsite II |  |  |  |  |
| --- | --- | --- | --- | --- | --- | --- | --- | --- | --- | --- |
|  | Ag0826 | T77 | Q140 | Y163 | L191 | G78 | T79 | A83 | H138 | F220 |
| I | LCC | Y95 | M166 | W190 | V212 | T96 | A97 | S101 | H164 | F243 |
| IIa | PE-H | F98 | M172 | W195 | I219 | V99 | S100 | S104 | W170 | Y250 |
| IIb | *Is*PETase | Y87 | M161 | W185 | I208 | T88 | A89 | S93 | W159 | S238 |
| III | *Halo*PETase1 | T88 | Q157 | Y180 | I208 | T89 | A90 | T94 | H155 | F237 |

## Acknowledgments

This work was supported by the Japan Society for the Promotion of Science (JSPS) through JSPS KAKENHI (25K15498 to S.H.) and the Environment Research and Technology Development Fund of the Environmental Restoration and Conservation Agency provided by Ministry of the Environment of Japan (JPMEERF20241RB1 to S.H.). We thank Ms. Aika Fujiki for her technical assistance. We also thank Open Facility Division, Global Facility Center, Creative Research Institution, Hokkaido University for performing NMR analysis and SEM analysis.

## Appendixes

The following is the Supplementary data to this article.

**Table S1.**
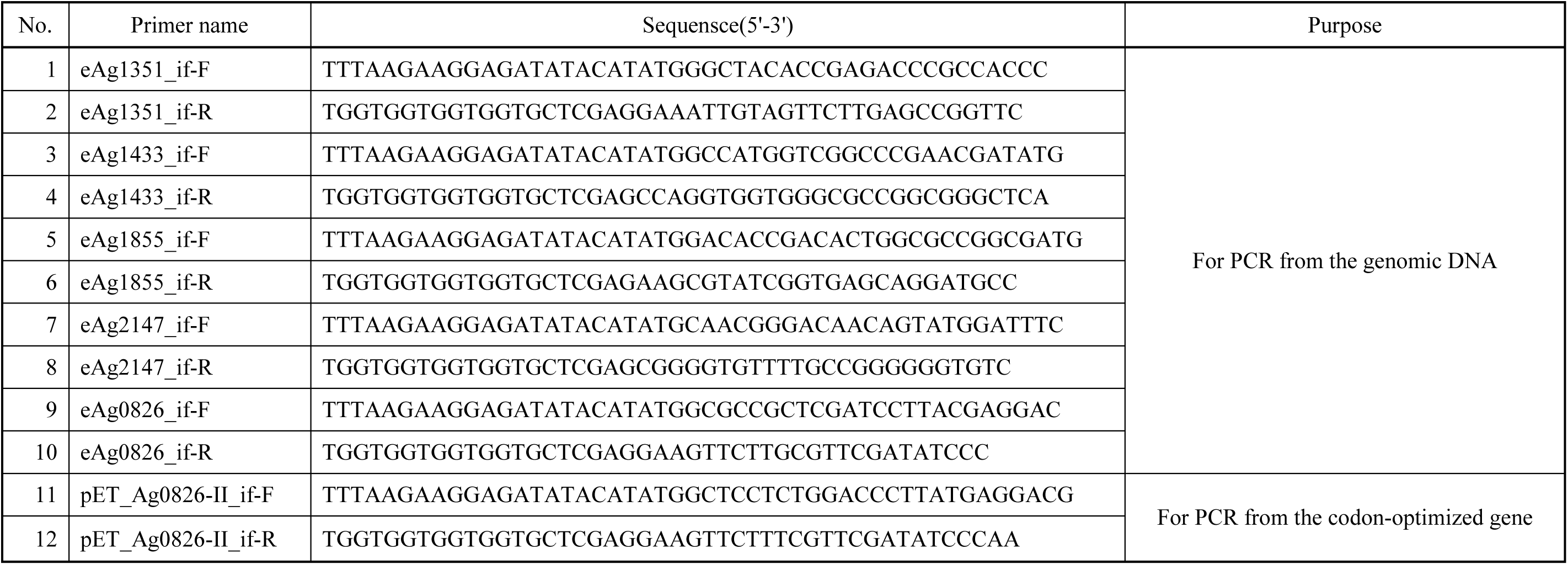
Primers used in this study.

**Fig. S1.**
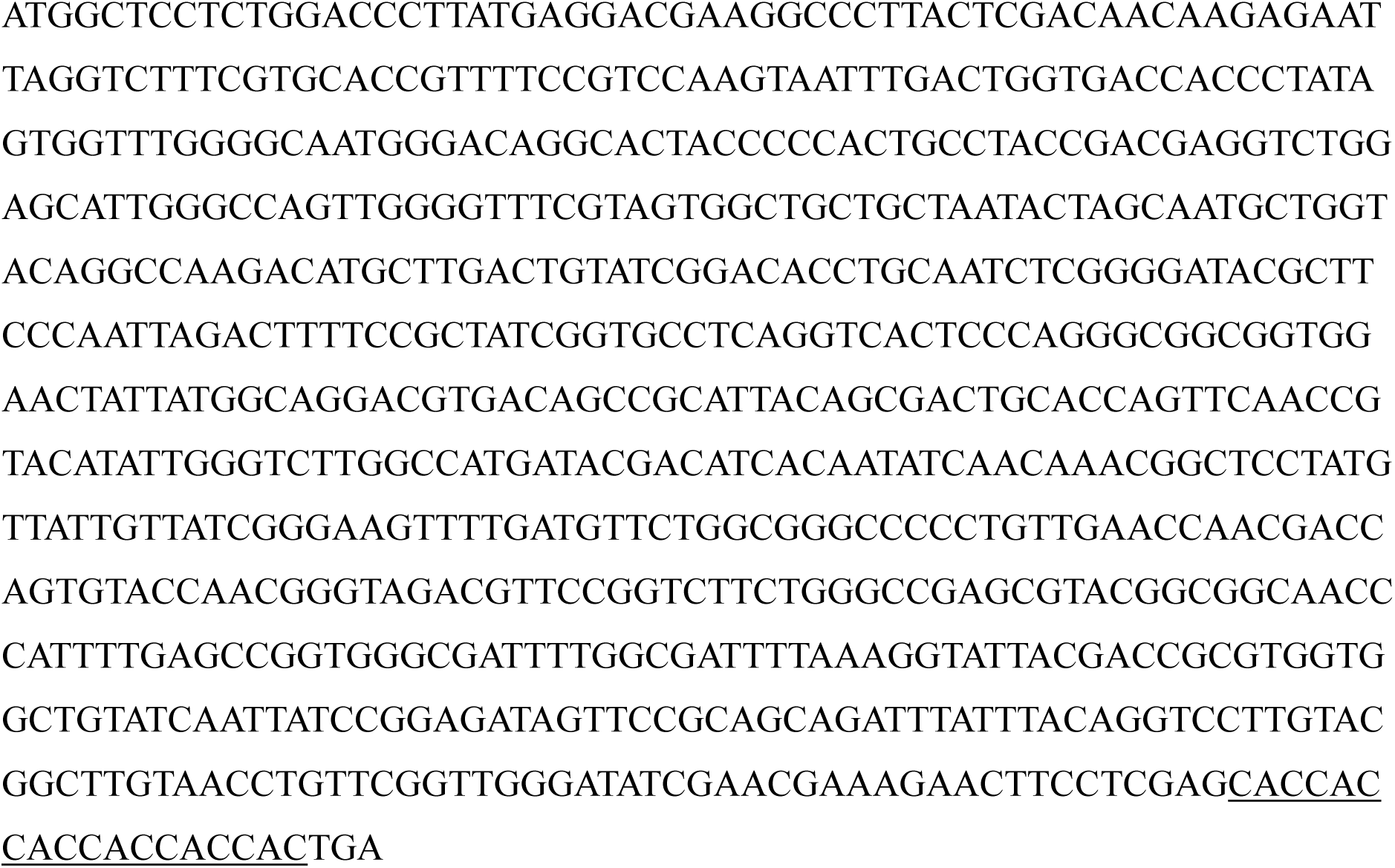
DNA sequence of the Ag0826(opt) coding region. The His_6_-tag is indicated by an underline.

**Fig. S2.**
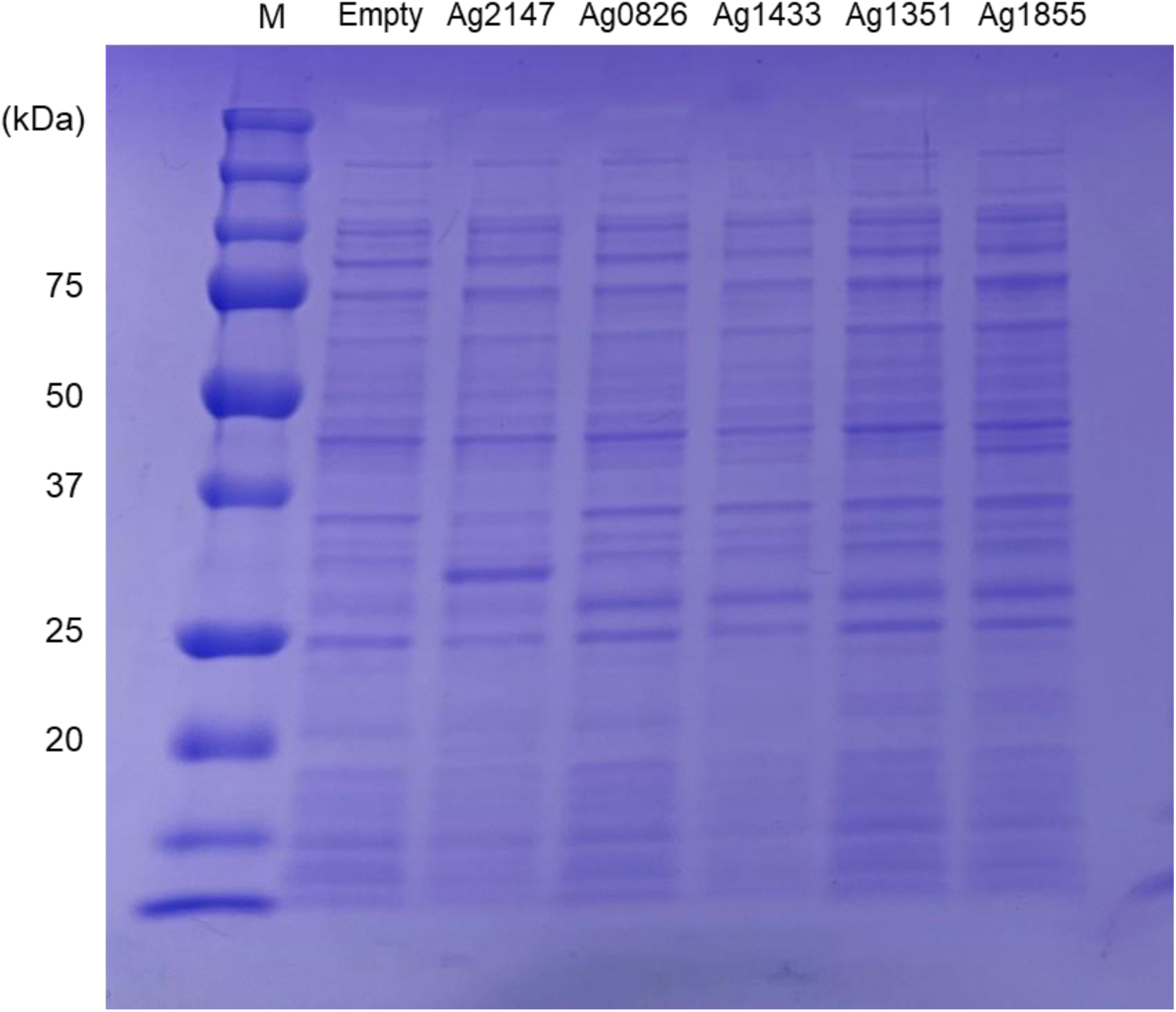
SDS-PAGE analysis of cell-free extracts expressing AgXXXX proteins. The “Empty” lane contains a cell-free extract from *E. coli* harboring the pET-21a(+) empty vector. Five micrograms of total protein was loaded per lane. Lane “M” indicates a molecular weight marker. The calculated molecular weights of the His_6_-tagged proteins are as follows: Ag2147, 33.11 kDa; Ag0826, 26.52 kDa; Ag1433, 62.08 kDa; Ag1351, 31.76 kDa; and Ag1855, 39.16 kDa.

**Fig. S3.**
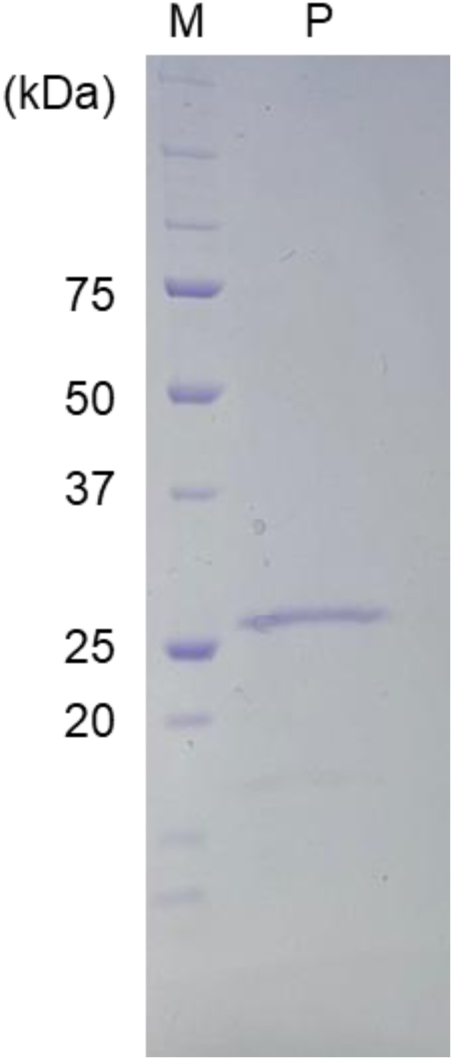
SDS-PAGE analysis of purified Ag0826. Lane “M” indicates a molecular weight marker, and lane “P” contains purified Ag0826. Five micrograms of protein were loaded in lane “P.”

**Fig. S4.**
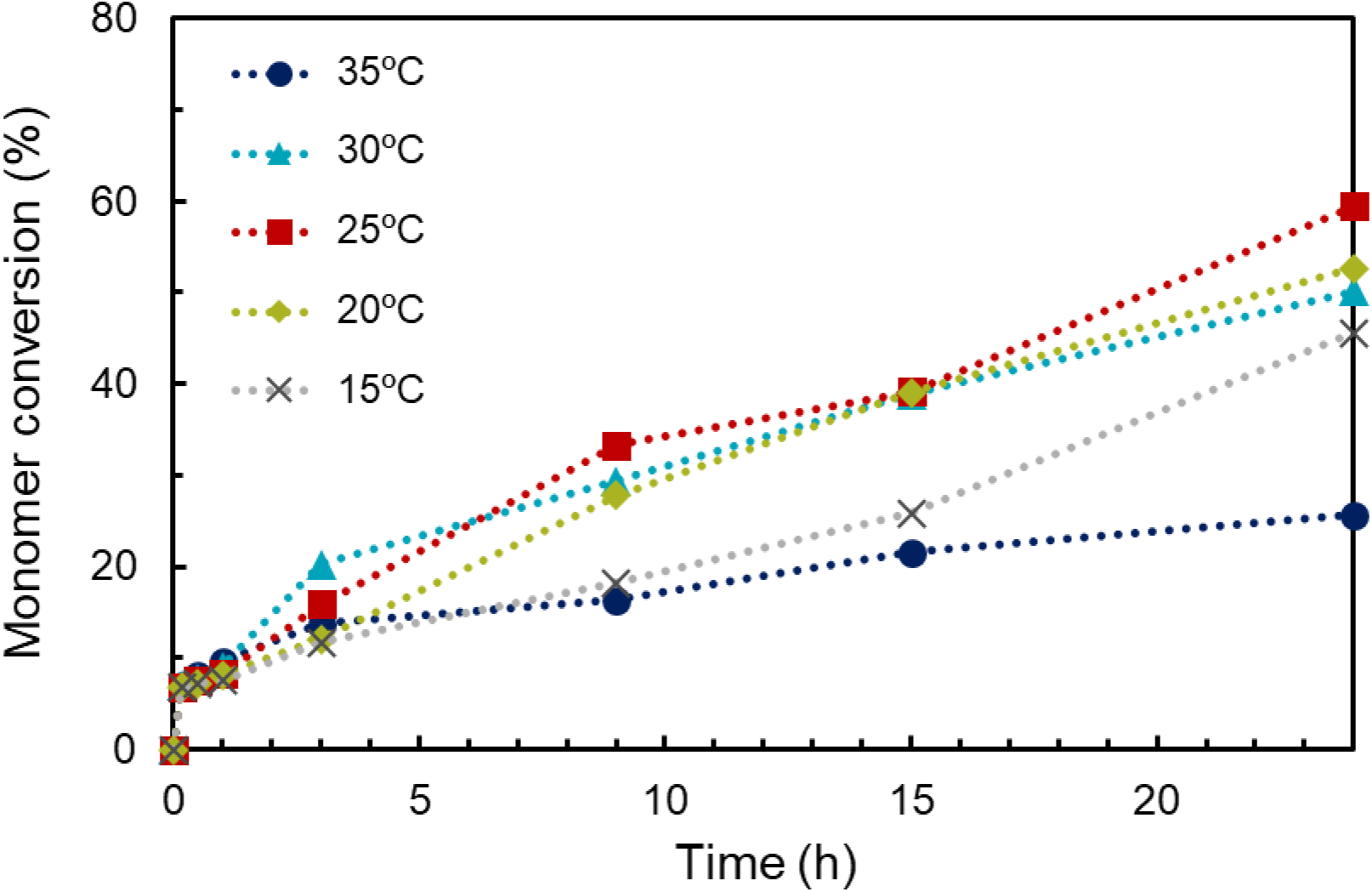
Time course of monomer (6HHx) production during PCL degradation by Ag0826 at different reaction temperatures. The reaction mixture consisted of 100 mM KPB (pH 8.0), 5.0 mg/mL PCL emulsion, and 2 µg/mL purified Ag0826, and the reaction was initiated by the addition of the enzyme. Reactions were incubated for up to 24 h at 15, 20, 25, 30, or 35°C.

**Fig. S5.**
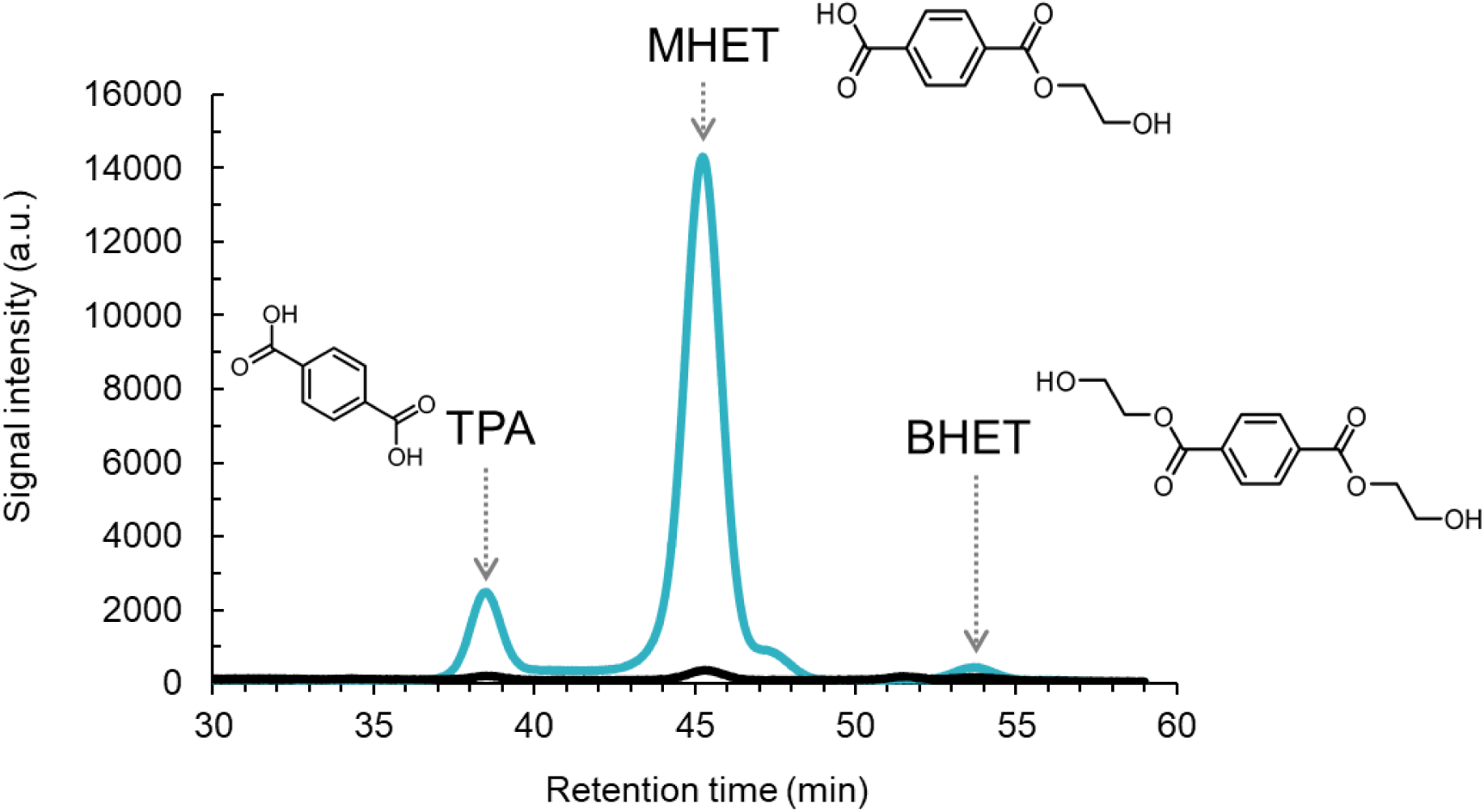
HPLC chromatograms of reaction supernatants after PET degradation assays monitored by UV detection at 240 nm. The light blue trace represents the reaction with Ag0826, and the black trace represents the reaction without enzyme. Peaks were identified by comparison with authentic TPA, MHET, and BHET standards.

